# A new highly sensitive real-time quantitative-PCR method for detection of *BCR-ABL1* to monitor minimal residual disease in chronic myeloid leukemia after discontinuation of imatinib

**DOI:** 10.1101/455956

**Authors:** Hiroaki Kitamura, Yoko Tabe, Koji Tsuchiya, Maiko Yuri, Tomohiko Ai, Shigeki Misawa, Takashi Horii, Akimichi Ohsaka, Shinya Kimura

**Author notes:** Corresponding author (Y.T.).

## Abstract

Tyrosine kinase inhibitors (TKIs) targeting the *BCR-ABL1* fusion protein, encoded by the Philadelphia chromosome, have drastically improved the outcomes for patients with chronic myeloid leukemia (CML). Although several real-time quantitative polymerase chain reaction (RQ-PCR) kits for the detection of *BCR-ABL1* transcripts are commercially available, their accuracy and efficiency in laboratory practice require reevaluation. We have developed a new in-house RQ-PCR method to detect minimal residual disease (MRD) in CML cases. MRD was analyzed in 102 patients with CML from the DOMEST study, a clinical trial to study the rationale for imatinib mesylate discontinuation in Japan. The *BCR-ABL1*/*ABL1* ratio was evaluated using the international standard (IS) ratio, where IS < 0.01% was defined as a major molecular response. At enrollment, *BCR-ABL1* transcripts were undetectable in all samples using a widely-applied RQ-PCR method performed in the commercial laboratory, BML (BML Inc., Tokyo, Japan); however, the in-house method detected the *BCR-ABL1* transcripts in five samples (5%) (mean IS ratio: 0.0062 ± 0.0010%). After discontinuation of imatinib, *BCR-ABL1* transcripts were detected using the in-house RQ-PCR in 21 patients (21%) that were not positive using the BML method. Nineteen samples were also tested using a commercially available RQ-PCR assay kit with a detection limit of IS ratio, 0.0007% (ODK-1201, Otsuka Pharmaceutical Co., Tokyo, Japan). This method detected low levels of *BCR-ABL1* transcripts in 14 samples (74%), but scored negative for five samples (26%) that were positive using the in-house method. These data suggest that our new in-house RQ-PCR method is effective for monitoring MRD in CML.

## Introduction

Chronic myeloid leukemia (CML) is a disease that arises in hematopoietic stem cells and is caused by a reciprocal translocation between chromosomes 9 and 22 (t(9;22)(q34;q11.2)), referred to as the Philadelphia chromosome, which generates *BCR-ABL1* fusion transcripts. The *BCR-ABL1* protein constitutively activates tyrosine kinase (TK) (1) that causes unregulated proliferation of abnormal blood cells, and consequently interrupts normal hematopoiesis. Theoretically, TK inhibition was expected to be an effective cure for CML, and imatinib, which competitively inhibits phosphorylation of BCR-ABL1, was developed in 2001 and is used as a frontline TK inhibitor (TKI) (2-5). Currently, according to the European Society of Medical Oncology (ESMO) Clinical Practice Guideline (2017), three commercially available TKIs, imatinib, dasatinib, and nilotinib, can be used for the CML therapy with no significant difference in survival rate (6).

To monitor the response to treatment with TKI, several assessment methods have been employed as follows: (1) complete hematologic response, determined by examination of complete blood cell counts and differentiated by flow cytometry; (2) complete cytogenetic response, evaluated using bone marrow aspirate and biopsy samples; and (3) molecular response (MR) examined by real-time quantitative-PCR (RQ-PCR) (7). Of these, MR detection by RQ-PCR is the most sensitive method to monitor minimal residual disease (MRD); however, RQ-PCR protocols vary among laboratories, potentially leading to inconsistencies in patient treatment. Therefore, the European Leukemia Network (ELN) and National Comprehensive Cancer Network (NCCN) have recommended monitoring *BCR-ABL1* mRNA levels by RQ-PCR using international standards (IS) (8-10). The International Randomized Study of Interferon versus STI571 (IRIS) proposed that log reduction of *BCR-ABL1*^IS^ (IS ratio) during therapy, compared with baseline IS ratio at diagnosis (*BCR-ABL1*^IS^, 100%), should be evaluated to monitor MRD. Initially, major molecular response (MMR), defined as *BCR-ABL1*^IS^ ≤ 0.1% (MR: 3.0; 3 log reduction) was considered adequate (11). Subsequently, deeper molecular responses (DMRs) were determined to be desirable. DMRs are defined as *BCR-ABL1*^IS^ ≤ 0.01% (MR: 4.0; 4 log reduction), *BCR-ABL1*^IS^ ≤ 0.0032% (MR: 4.5; 4.5 log reduction), and *BCR-ABL1*^IS^ ≤ 0.001% (MR: 5.0; 5 log reduction) (12).

Long-term treatment with TKIs can cause considerable adverse effects, including gastrointestinal damage, fluid retention, bone marrow suppression, liver injury, cardiovascular events, and kidney injury (13). Even more seriously, some patients develop resistance to imatinib (14). In some cases, imatinib must be discontinued or replaced with a different TKI, such as bosutinib, because of such problems. Consequently, several clinical trials have been conducted to investigate whether TKIs can be ceased after DMR is achieved. Mahon *et al*. reported that approximately 40% of patients with CML remained in complete molecular response (CMR) for at least 2 years after discontinuation of imatinib (15). Stop studies of second-generation TKIs (dasatinib and nilotinib) showed that approximately 50% of patients achieved, and remained in, DMR following TKI cessation (16, 17). As DMR is an emerging goal in CML and necessary for entry into treatment discontinuation studies (15, 18, 19), RQ-PCR assays with inadequate sensitivity could fail to detect low level *BCR-ABL1* fusion transcripts, leading to inappropriate or premature treatment cessation attempts. Therefore, well defined guidelines have been developed to ensure adequate sensitivity levels are achieved, down to MR4.0 or MR4.5 (20). The World Health Organization International Genetic Reference Panel for the quantitation of *BCR-ABL1* mRNA (World Health Organization document, World Health Organization/BS/09.2106) has been distributed to manufacturers to generate secondary reference materials (21), and commercial kits are now available from several manufacturers (22).

Recently, we developed a new in-house RQ-PCR method and determined its sensitivity as 0.0033% using synthetic ARQ IS Calibrator Panels; this level of sensitivity is sufficient to detect MRD (23). In this study, we evaluated the ability of this in-house RQ-PCR method to detect low level *BCR-ABL1* fusion transcripts using samples obtained in the ongoing Delightedly Overcome CML Expert Stop TKI (DOMEST) clinical trial to evaluate the rationale for cessation of imatinib (24).

## Materials and Methods

### Study design

This study was performed as a part of the DOMEST clinical trial, which is being conducted to elucidate the rationale for imatinib discontinuation in Japan. The enrollment criteria were (1) 15 years of age or older, (2) diagnosed with CML in chronic phase and receiving imatinib therapy, and (3) maintained DMR for longer than 2 years (MR4.0 or MR4.0 equivalent), as determined by transcription-mediated amplification, reverse transcriptase-polymerase chain reaction (RT-PCR), or real-time quantitative polymerase chain reaction (RQ-PCR). Other inclusion criteria were a WHO performance status score of 0–2 and absence of severe dysfunction of primary organs. Previous therapies additional to imatinib were permitted. Patients with additional chromosomal abnormalities and those with a positive RQ-PCR result using the method applied by BML (BML Inc., Tokyo, Japan) at the time of registration were excluded. The study was approved by the ethics committees of Saga University Graduate School of Medicine and Juntendo University Graduate School of Medicine. All participants provided written informed consent for their samples and data from their medical records to be used for research.

In the DOMEST study, RQ-PCR was performed every month for the first year and every 3 months for the second year by BML (16, 25); molecular recurrence was defined as *BCR-ABL1* detected by two successive tests, or by loss of MR3.0 in one test. Residual total RNA samples from the study were subsequently used for measurement using the in-house RQ-PCR method. The major *BCR-ABL1* mRNA assay kit, ODK-1201 (Otsuka Pharmaceutical Co., Japan), which also uses the RQ-PCR technique, was used to test available samples showing discrepant results between the in-house and BML methods for comparison (26).

### RNA extraction and cDNA synthesis

Total RNA was extracted from 7 mL peripheral blood in EDTA tubes using a QIAamp RNA Blood Mini Kit (Qiagen, Hilden, Germany). RNA was quantified by Nanodrop spectrophotometry (ND 2000-NanoDrop 3.2.1, Thermo Scientific, Waltham, USA). Transcriptor Universal cDNA Master reverse transcriptase (Roche Diagnostics, Mannheim, Germany) was used for cDNA synthesis, using 1 µg total RNA.

### Quantitative real-time PCR

cDNA was amplified by 55 cycles of RT-PCR in a final reaction volume of 20 µL using the LightCycler^®^ 2.0 (Roche Diagnostics, Mannheim, Germany) and LightCycler^®^ TaqMan^®^ Master, in accordance with the manufacturer’s instructions. *ABL1* was used as the control gene. The primers and probes used were as follows: *BCR-ABL1* forward primer, 5′-TGACCAACTCGTGTGTGAAACTC-3′, reverse primer, 5′- CACTCAGACCCTGAGGCTCAA-3′, and probe, 5′-CCCTTCAGCGGCCAGTAGCATCTGA-3′; *ABL1* forward primer, 5′-CGAAGGGAGGGTGTACCATTA-3′, reverse primer, 5′- CAACTCGGCCAGGGTGTT-3′, and probe, 5′- CTTCTGATGGCAAGCTCTACGTCTCCTCC-3′. Sequences were obtained from GenBank (Accession Nos. X02596 for *BCR* and X16416 for *ABL1*). Probes contained the fluorescent reporter dye, 6-carboxyfluorescein (FAM), at the 5′-end and the fluorescent quencher dye, Black Hole Quencher (BHQ), at the 3′-end. Results are reported as *BCR-ABL1*/*ABL1* ratios (%).

### RNA standards for the RQ-PCR assay

An *in vitro* transcribed RNA from the *BCR-ABL1* gene of the K562 cell line was used to determine the lower detection limit of the assay. A region of 188 bp, including the *BCR-ABL1* breakpoint, was amplified by PCR using the primers described above. The product was purified using the QIAquick PCR Purification Kit^TM^ (Qiagen, Hilden, Germany) and then ligated to the pGEM-T vector (Promega, Madison, USA). The recombinant plasmid was transformed to the DH5α *Escherichia coli* strain (Promega), and the cloned plasmid was extracted using a QIAprep Spin Miniprep Kit^TM^ (Qiagen). The orientation of the DNA insert was confirmed by sequencing. *In vitro* transcription was performed using either the RiboMAX Large Scale RNA Production System^TM^ or the T7 RiboMAX Express Large Scale RNA Production System^TM^ (Promega), depending on the direction of inserts, as determined by sequencing. Transcribed RNA was purified using the RNeasy Mini Kit^TM^ (Qiagen), and the amount of RNA was quantified using the Agilent RNA 6000 Nano Assay^TM^ (Agilent Technology, California, USA).

The RNA copy number (/μl) was calculated using the following equation:

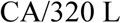

where C is the concentration of RNA (g/μl), assessed using the Agilent 2100 Bioanalyzer; A is Avogadro’s constant (6.0 × 10^23^ copies/mol); L is the length of synthetic RNA (nucleotides); and 320 is an approximation of the molecular weight of a nucleotide (g/mol).

### Determination of a laboratory-specific correlation parameter (CP) and data analyses

The World Health Organization (WHO) established an international genetic reference panel for quantification of *BCR-ABL1* fusion transcripts by RQ-PCR, which contains four different ratios (10%, 1%, 0.1%, and 0.01%) using the *BCR-ABL1*-positive cell line, K562, diluted in the *BCR-ABL1*-negative cell line, HL60 (21). Four level Armored RNA Quant (ARQ) (Asuragen, Inc., Austin, TX, USA) secondary reference panels were manufactured based on the WHO primary standards (22). Laboratory-specific CP equivalent conversion factor values were calculated for use with the ARQ IS Calibrator Panels.

The CP of the in-house RQ-PCR method was determined using the ARQ IS Calibrator Panel^TM^ containing four calibrators: IS 4.1%, 0.37%, 0.027%, and 0.0033%. The CP value for this study was 18.39. The *BCR-ABL1* mRNA ratio of standard material RNAs supplied by the panel was quantified using the local method, and 95% limits of agreement (LOA) were calculated. Values outside of the 95% LOA were omitted, and the CP was calculated by dividing the measured value by the expected value.

### Sequencing analysis

*BCR-ABL1* PCR products were separated and purified using agarose gel electrophoresis and a QIAquick Gel Extraction Kit (Qiagen). Cycle sequencing was performed using a BigDye Terminator v1.1 Cycle Sequencing Kit (Applied Biosystems, Foster City, USA). Cycle sequencing products were purified using a BigDye Xterminator Purification Kit (Applied Biosystems, Foster City, USA) before being run on an automated ABI 3500 genetic analyzer (Applied Biosystems), and sequences were analyzed using Sequencing Analysis software ver.6.

## Results

### Background of patients

Between January 2014 and May 2015, a total of 110 patients were enrolled for the DOMEST study; 104 of them were evaluated in this study. Among these patients, 102 were confirmed as having DMR (MR4.0) status, defined as “*BCR-ABL1* transcript levels below the detection limit of the widely-used BML method.” After MR4.0 (Log4) was confirmed, imatinib was ceased. The other two patients were excluded from this study because *BCR-ABL1* transcripts were detected at enrollment, and dasatinib was started. Patient characteristics are summarized in Table 1.

**Table 1.**
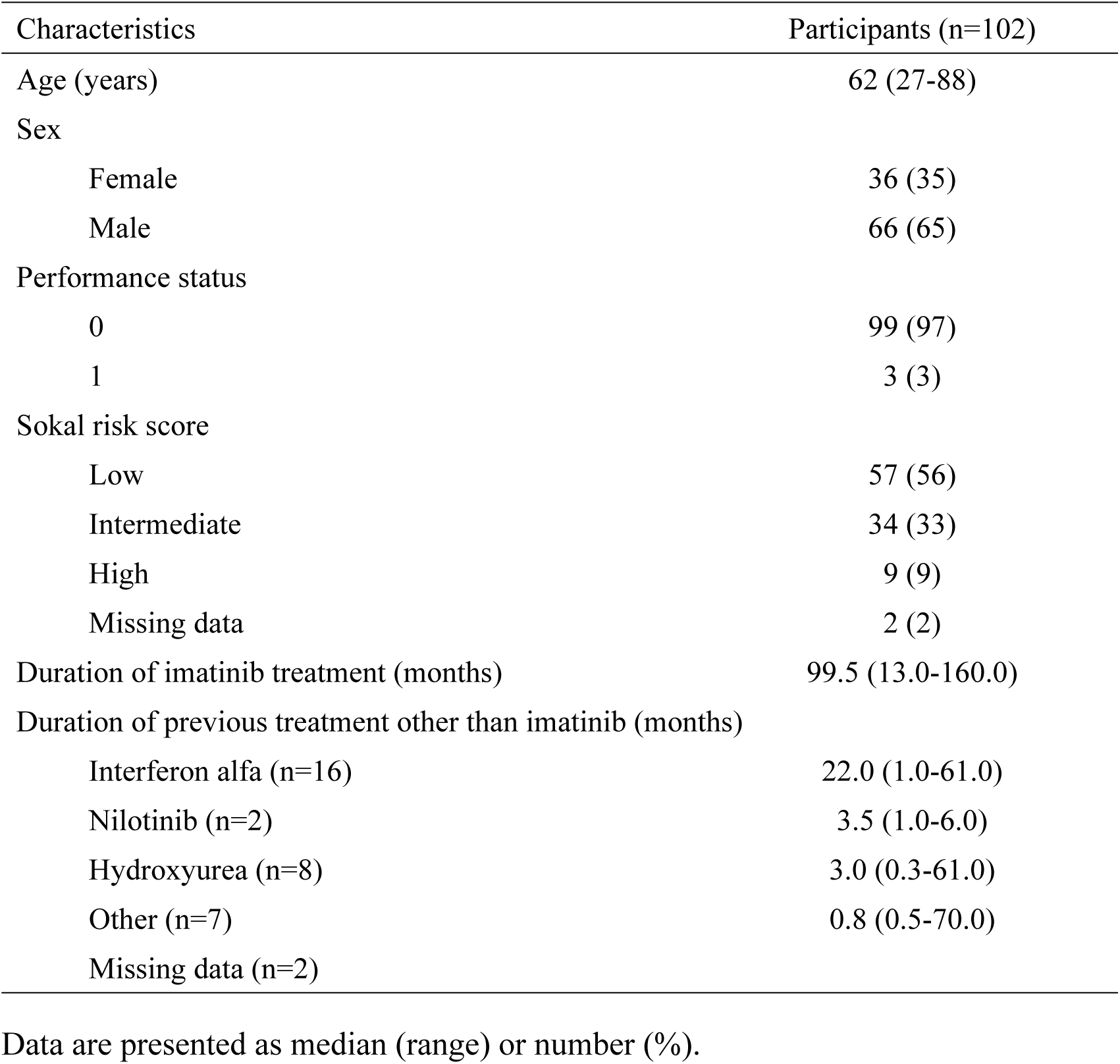
Patient characteristics

### Comparison of *BCR-ABL1* mRNA levels measured by the BML and in-house RQ-PCR methods during follow-up of imatinib discontinuation

Surprisingly, despite confirmation of DMR using the BML method in all enrolled cases, *BCR-ABL1* transcripts were detected by the in-house RQ-PCR method in 5 of 102 patients (5%) at the beginning of the DOMEST study. The IS ratios detected using the in-house method in these five cases are presented in S1 Table. The sequences of the PCR amplicons were confirmed by Sanger sequencing (S1 Fig); however, in the DOMEST study, these five patients remained in MR4.0, as determined by the BML method, throughout the study. Subsequently, *BCR-ABL1* fusion transcripts were detected in 15 cases (15%) by the BML method and the in-house method at the same time points, at an average ± standard deviation of 2.47 ± 2.13 months after cessation of imatinib (concordant cases, Table 2). In one case (patient #15), the fusion transcript level was < 0.01% by the BML method. In these recurred cases, TKI therapies were restarted in the DOMEST study.

**Table 2.**
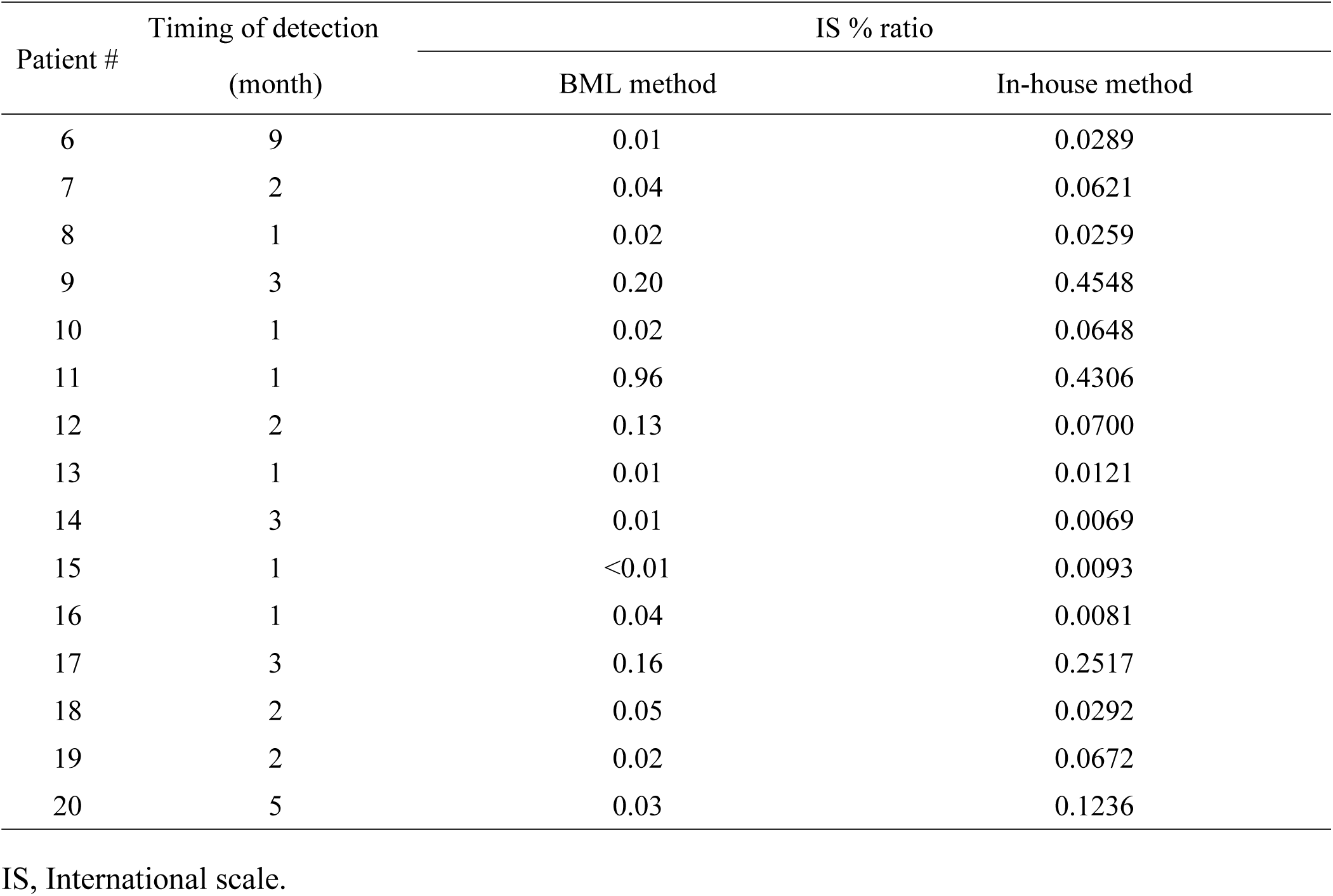
Cases with concordant results for detection of *BCR-ABL1* fusion transcripts using the in-house and BML methods

Amplicon sequences were confirmed by Sanger sequencing. Red arrows indicate the breakpoint (gcgg/ccag) of the *BCR-ABL1* fusion transcripts.

By contrast, the results were discordant between the in-house and BML methods in 21 cases (21%) (Table 3). The in-house method detected *BCR-ABL1* fusion transcripts at an average (range) of 2.4 (1–13) months earlier than the BML method. TKI therapies were restarted in these cases (Table 3). Of these 21 cases, 19 available samples were also tested using another RQ-PCR assay kit (ODK-1201; Otsuka Pharmaceutical, Japan), and the results were compared with those from the in-house method. As shown in Table 3, the ODK-1201 method detected a low level (IS < 0.01%) of *BCR-ABL1* fusion transcripts in 14 samples (74%), while they were detected in all 19 cases using the in-house method.

**Table 3.** Cases with discordant results for detection of *BCR-ABL1* fusion transcripts using the in-house and BML methods

| Patient # | Time of detection (month) |  | IS ratio (%) |  |
| --- | --- | --- | --- | --- |
|  | BML | In-house | In-house method | ODK-1201 |
| 21 | 15 | 12 | 0.0201 | ND |
| 22 | 18 | 5 | 0.0041 | ND |
| 23 | 1 | 0 | 0.0081 | NA |
| 24 | 11 | 4 | 0.0184 | 0.0124 |
| 25 | 5 | 3 | 0.0584 | 0.0107 |
| 26 | 1 | 0 | 0.0326 | 0.0021 |
| 27 | 4 | 2 | 0.0269 | 0.0061 |
| 28 | 15 | 12 | 0.0141 | 0.0054 |
| 29 | 2 | 1 | 0.0115 | 0.0040 |
| 30 | 1 | 0 | 0.0127 | NA |
| 31 | 3 | 2 | 0.0284 | 0.0186 |
| 32 | 1 | 0 | 0.0120 | ND |
| 33 | 5 | 4 | 0.0353 | 0.0214 |
| 34 | 4 | 3 | 0.0481 | 0.0039 |
| 35 | 2 | 0 | 0.0069 | 0.0024 |
| 36 | 1 | 0 | 0.0194 | 0.0054 |
| 37 | 6 | 3 | 0.0010 | 0.0049 |
| 38 | 3 | 0 | 0.0082 | ND |
| 39 | 1 | 0 | 0.0175 | 0.0058 |
| 40 | 1 | 0 | 0.0084 | 0.0045 |
| 41 | 3 | 1 | 0.0019 | ND |

ND, not detected; NA, not applicable.

*BCR-ABL1* fusion transcripts were detected by the in-house method in all samples positive by the BML method. In the remaining 61 cases, *BCR-ABL1* fusion transcripts were not detected using either the in-house or BML methods throughout the study.

## Discussion

Detection of low levels of *BCR-ABL1* fusion transcripts to monitor MRD must be performed quickly, efficiently, and at a reasonable cost (27). Currently, several commercial kits using IS are available (22, 28); however, these are designed to test large-scale samples in commercial laboratories, rather than for hospital laboratories, in which relatively small numbers of samples are examined. Therefore, we developed an in-house RQ-PCR method that is more accurate and flexible than those currently in use (23). In the present study, we evaluated the relevance and accuracy of this in-house RQ-PCR method using samples obtained for the DOMEST trial. Our data demonstrate that the in-house method is sufficiently sensitive to detect MRD and recurrence, relative to the widely-used BML method and the recently developed ODK-1201 commercial kit.

In the DOMEST trail, clinical decisions were made based on the monitoring of *BCR-ABL1* fusion transcripts measured using the BML method (detection limit: IS 0.01%) (BML Inc.) (25). The *BCR-ABL1* mRNA quantification results obtained using the in-house RQ-PCR agreed with those generated using the BML method in 15 cases (Table 2). By contrast, the in-house method detected MRD earlier than the BML method in 21.0% of cases (Table 3). Although both the in-house and ODK-1201 methods can identify at least a 4.5 log reduction in the IS ratio (26), the IS ratios measured in this study were somewhat discordant between the two methods. In five samples, the IS ratios were below the detection limit (IS ratio < 0.0007%) of the ODK-1201 method, whereas using the in-house method they had a mean IS ratio of 0.0094 ± 0.00754%. In 13 cases, the IS ratios determined using the ODK-1201 method were lower than those using the in-house method. This discordance may be because of RNA degradation, since measurements could not all be performed at exactly the same time. Alternatively, it could be due to the relatively large variability in the detection of very low copy number transcripts, which is unavoidable using current technology.

In recent clinical trials to evaluate the rationale for TKI cessation, the sensitivity of assays used for detection of *BCR-ABL1* fusion transcripts has been claimed as 5 logs (12, 29); however, there are reasons to be skeptical about the accuracy of measurements of such extremely low amounts of mRNA. Even a subtle pipetting error can easily lead to an enormous difference. In addition, despite using the IS, calculation and/or methods of determining conversion factors can significantly affect the results. According to the UKNEQAS (external quality assessment), the variability of results among participant laboratories was considerable, even after the introduction of an IS (30). Therefore, the methodology used for the measurement of *BCR-ABL1* fusion transcripts requires further improvement.

This study has certain limitations. The clinical relevance of our new in-house RQ-PCR is uncertain since it has not been assessed in large-scale randomized clinical trials. Moreover, very long-term outcomes of imatinib therapy in CML have yet to be elucidated (31). Current recommendations for the definition of MRs may be changed after accumulation of further data. Although we performed all experimental procedures with great care, the introduction of some errors caused by human factors cannot be completely excluded, as noted in a recent commentary (32).

In conclusion, our newly developed in-house RQ-PCR method with IS calibration was accurate and effective for detecting MRD in the context of an imatinib cessation study. The main advantages of this assay lie in the promptness with which results are obtained and its ease of use. Thus, this method could be advantageous for implementation in hospital laboratories, where small numbers of samples are tested.

## Acknowledgments

The authors would like to thank all the patients and physicians who participated in this study. We thank Yumi Miyashita (ECRIN, Japan) for monitoring the clinical trial, and Makoto Nagano (BML) and Hiromi Yamada for technical support.

## Supporting information

**S1_Fig.**
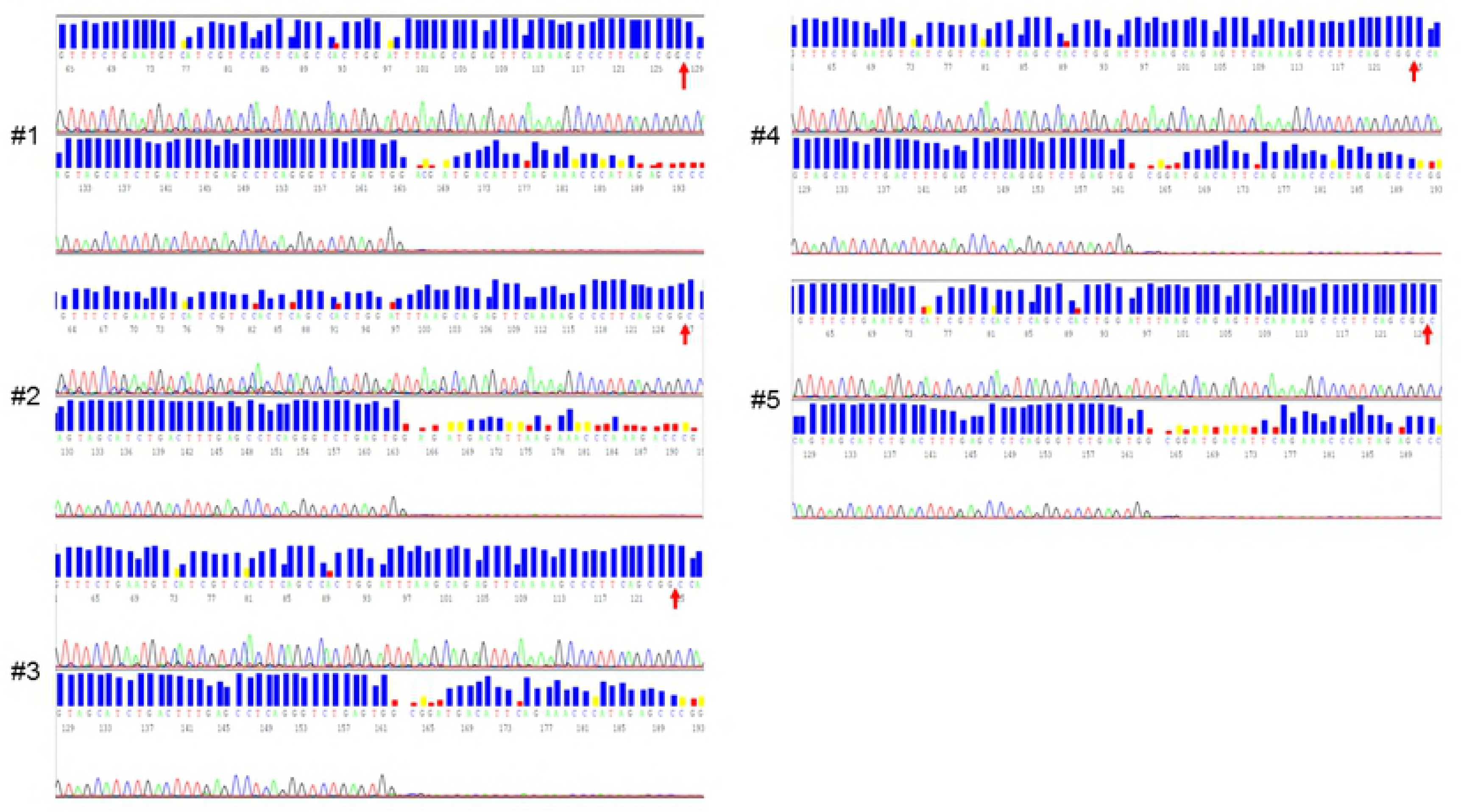
**PCR amplicon sequence data for patients detected by in-house, but not BML, methods at the beginning of the study.**

**S1 Table.** **IS ratios generated using the in-house RQ-PCR method for samples where transcripts were not detected by the BML method at the beginning of the study.**

| Patient # | IS % ratio (In-house method) |
| --- | --- |
| 1 | 0.0077 |
| 2 | 0.0057 |
| 3 | 0.0067 |
| 4 | 0.0052 |
| 5 | 0.0059 |

## References

1. Kimura S, Ando T, Kojima K. Ever-advancing chronic myeloid leukemia treatment. Int J Clin Oncol. 2014;19(1):3-9.

2. O’Brien SG, Guilhot F, Larson RA, Gathmann I, Baccarani M, Cervantes F, et al. Imatinib compared with interferon and low-dose cytarabine for newly diagnosed chronic-phase chronic myeloid leukemia. N Engl J Med. 2003;348(11):994-1004.

3. Saglio G, Kim DW, Issaragrisil S, le Coutre P, Etienne G, Lobo C, et al. Nilotinib versus imatinib for newly diagnosed chronic myeloid leukemia. N Engl J Med. 2010;362(24):2251-9.

4. Kantarjian H, Shah NP, Hochhaus A, Cortes J, Shah S, Ayala M, et al. Dasatinib versus imatinib in newly diagnosed chronic-phase chronic myeloid leukemia. N Engl J Med. 2010;362(24):2260-70.

5. Cortes JE, Kim DW, Kantarjian HM, Brummendorf TH, Dyagil I, Griskevicius L, et al. Bosutinib versus imatinib in newly diagnosed chronic-phase chronic myeloid leukemia: results from the BELA trial. J Clin Oncol. 2012;30(28):3486-92.

6. Hochhaus A, Saussele S, Rosti G, Mahon FX, Janssen J, Hjorth-Hansen H, et al. Chronic myeloid leukaemia: ESMO Clinical Practice Guidelines for diagnosis, treatment and follow-up. Ann Oncol. 2017;28(suppl_4):iv41-iv51.

7. Cortes J, Rousselot P, Kim DW, Ritchie E, Hamerschlak N, Coutre S, et al. Dasatinib induces complete hematologic and cytogenetic responses in patients with imatinib-resistant or -intolerant chronic myeloid leukemia in blast crisis. Blood. 2007;109(8):3207-13.

8. Baccarani M, Deininger MW, Rosti G, Hochhaus A, Soverini S, Apperley JF, et al. European LeukemiaNet recommendations for the management of chronic myeloid leukemia: 2013. Blood. 2013;122(6):872-84.

9. Network NCC. National Comprehensive Cancer Network - Chronic Myloid Leukemia (Version 3.2018) Available from: www.nccn.org.

10. Baccarani M, Cortes J, Pane F, Niederwieser D, Saglio G, Apperley J, et al. Chronic myeloid leukemia: an update of concepts and management recommendations of European LeukemiaNet. J Clin Oncol. 2009;27(35):6041-51.

11. Hughes TP, Kaeda J, Branford S, Rudzki Z, Hochhaus A, Hensley ML, et al. Frequency of major molecular responses to imatinib or interferon alfa plus cytarabine in newly diagnosed chronic myeloid leukemia. N Engl J Med. 2003;349(15):1423-32.

12. Cross NC, White HE, Muller MC, Saglio G, Hochhaus A. Standardized definitions of molecular response in chronic myeloid leukemia. Leukemia. 2012;26(10):2172-5.

13. Bauer S, Buchanan S, Ryan I. Tyrosine Kinase Inhibitors for the Treatment of Chronic-Phase Chronic Myeloid Leukemia: Long-Term Patient Care and Management. J Adv Pract Oncol. 2016;7(1):42-54.

14. Rossari F, Minutolo F, Orciuolo E. Past, present, and future of Bcr-Abl inhibitors: from chemical development to clinical efficacy. J Hematol Oncol. 2018;11(1):84.

15. Mahon FX, Rea D, Guilhot J, Guilhot F, Huguet F, Nicolini F, et al. Discontinuation of imatinib in patients with chronic myeloid leukaemia who have maintained complete molecular remission for at least 2 years: the prospective, multicentre Stop Imatinib (STIM) trial. Lancet Oncol. 2010;11(11):1029-35.

16. Imagawa J, Tanaka H, Okada M, Nakamae H, Hino M, Murai K, et al. Discontinuation of dasatinib in patients with chronic myeloid leukaemia who have maintained deep molecular response for longer than 1 year (DADI trial): a multicentre phase 2 trial. Lancet Haematol. 2015;2(12):e528-35.

17. Rea D, Nicolini FE, Tulliez M, Guilhot F, Guilhot J, Guerci-Bresler A, et al. Discontinuation of dasatinib or nilotinib in chronic myeloid leukemia: interim analysis of the STOP 2G-TKI study. Blood. 2017;129(7):846-54.

18. Ross DM, Hughes TP. How I determine if and when to recommend stopping tyrosine kinase inhibitor treatment for chronic myeloid leukaemia. Br J Haematol. 2014;166(1):3-11.

19. Takahashi N, Tauchi T, Kitamura K, Miyamura K, Saburi Y, Hatta Y, et al. Deeper molecular response is a predictive factor for treatment-free remission after imatinib discontinuation in patients with chronic phase chronic myeloid leukemia: the JALSG-STIM213 study. Int J Hematol. 2017.

20. Cross NC, White HE, Colomer D, Ehrencrona H, Foroni L, Gottardi E, et al. Laboratory recommendations for scoring deep molecular responses following treatment for chronic myeloid leukemia. Leukemia. 2015;29(5):999-1003.

21. White HE, Matejtschuk P, Rigsby P, Gabert J, Lin F, Lynn Wang Y, et al. Establishment of the first World Health Organization International Genetic Reference Panel for quantitation of BCR-ABL mRNA. Blood. 2010;116(22):e111-7.

22. White HE, Hedges J, Bendit I, Branford S, Colomer D, Hochhaus A, et al. Establishment and validation of analytical reference panels for the standardization of quantitative BCR-ABL1 measurements on the international scale. Clin Chem. 2013;59(6):938-48.

23. Yamada H, Tabe Y, Watanabe K, Morishita S, Yuri M, Yokoo M, et al. Harmonization of quantitative BCR-ABL measurements using the secondary reference material anchored to the WHO primary standards. Int J Lab Hematol. 2015;37(2):e29-33.

24. Fujisawa S, Ueda Y, Usuki K, Kobayashi H, Kondo E, Doki N, et al. Feasibility of the Imatinib Stop Study in the Japanese Clinical Setting: Delightedly Overcome CML Expert Stop TKI Trial (DOMEST Trial). Int J Clin Oncol. revising.

25. Yoshida C, Fletcher L, Ohashi K, Wakita H, Kumagai T, Shiseki M, et al. Harmonization of molecular monitoring of chronic myeloid leukemia therapy in Japan. Int J Clin Oncol. 2012;17(6):584-9.

26. Nakamae H, Yoshida C, Miyata Y, Hidaka M, Uike N, Koga D, et al. A new diagnostic kit, ODK-1201, for the quantitation of low major BCR-ABL mRNA level in chronic myeloid leukemia: correlation of quantitation with major BCR-ABL mRNA kits. Int J Hematol. 2015;102(3):304-11.

27. Arber DA, Borowitz MJ, Cessna M, Etzell J, Foucar K, Hasserjian RP, et al. Initial Diagnostic Workup of Acute Leukemia: Guideline From the College of American Pathologists and the American Society of Hematology. Arch Pathol Lab Med. 2017;141(10):1342-93.

28. Maute C, Nibourel O, Rea D, Coiteux V, Grardel N, Preudhomme C, et al. Calibration of BCR-ABL1 mRNA quantification methods using genetic reference materials is a valid strategy to report results on the international scale. Clin Biochem. 2014;47(13-14):1333-6.

29. Laneuville P. When to Stop Tyrosine Kinase Inhibitors for the Treatment of Chronic Myeloid Leukemia. Curr Treat Options Oncol. 2018;19(3):15.

30. Scott S, Travis D, Whitby L, Bainbridge J, Cross NCP, Barnett D. Measurement of BCR-ABL1 by RT-qPCR in chronic myeloid leukaemia: findings from an International EQA Programme. Br J Haematol. 2017;177(3):414-22.

31. Nicolini FE, Alcazer V, Cony-Makhoul P, Heiblig M, Morisset S, Fossard G, et al. Long-term follow-up of de novo chronic phase chronic myelogenous leukemia patients on front-line imatinib. Exp Hematol. 2018.

32. Arora R, Press RD. Measurement of BCR-ABL1 transcripts on the International Scale in the United States: current status and best practices. Leuk Lymphoma. 2017;58(1):8-16.

